# A nepenthesin insert allosterically controls catalysis in the malaria parasite protease plasmepsin V

**DOI:** 10.1101/2022.02.28.482356

**Authors:** Alexander J. Polino, Justin J. Miller, Soumendranath Bhakat, Suhas Bobba, Gregory R. Bowman, Daniel E. Goldberg

**Author notes:** Address correspondence to Daniel E. Goldberg.

## Abstract

Plasmepsin V (PM V) is a pepsin-like aspartic protease essential for growth of the malaria parasite *Plasmodium falciparum*. Previous work has shown PM V to be an ER-resident protease that processes parasite proteins destined for export into the host cell. Depletion or inhibition of the enzyme is lethal during asexual replication within red blood cells, as well as during the formation of sexual stage gametocytes. The structure of the *P. vivax* PM V has been characterized by x-ray crystallography, revealing a canonical pepsin fold punctuated by structural features uncommon to secretory aspartic proteases. Here we use parasite genetics to probe these structural features by attempting to rescue lethal PM V depletion with various mutant enzymes. We find an unusual nepenthesin 1-type insert to be essential for parasite growth and PM V activity. Mutagenesis of the nepenthesin insert suggests that both its amino acid sequence and one of the two disulfide bonds that undergird its structure are required for the nepenthesin insert’s role in PM V function. Molecular dynamics simulations paired with Markov state modelling suggest that the nepenthesin insert allosterically controls PM V catalysis through multiple mechanisms.

## Introduction

Malaria remains a substantial burden on the developing world, with over 240 million cases and 600,000 deaths annually (1). While the parasite life cycle includes several host tissues, symptomatic human disease results entirely from infection of red blood cells (RBCs). After invasion of an RBC, the parasite dramatically modifies the host cell by exporting hundreds of effector proteins into the host cytosol. This program of protein export results in altered host cell rigidity, increased permeability to nutrients, and the constitution of a secretory apparatus that facilitates the display of parasite-encoded adhesins on the RBC surface (2, 3). These adhesins mediate binding of parasitized RBCs to host vascular endothelia, resulting in vascular disruption and, in extreme cases, death of the host.

Most known exported *P. falciparum* effectors bear a motif termed the *Plasmodium* export element (PEXEL) near the protein N-terminus (4, 5). PEXEL – with consensus sequence RxLxE/Q/D – is cleaved in the parasite ER by the aspartic protease plasmepsin V (PM V) (6, 7). Inhibition of PM V activity disrupts protein export and kills parasites (8). PM V is essential at two distinct points of intraerythrocytic development (9), as well as during gametocytogenesis (10), raising the profile of this enzyme as a potential target for chemotherapeutic intervention.

A recent x-ray crystallographic structure of the *P. vivax* PM V bound to the peptidomimetic inhibitor WEHI-842 revealed features unusual for a secretory aspartic protease (11). A nepenthesin 1-type insert and a helix-turn-helix domain feature prominently at opposite surfaces of the protein. The nepenthesin insert – so-called as it was identified in the protease nepenthesin 1 from the digestive fluid of the pitcher plant *Nepenthesia* – is a tri-lobed structure held together by two cysteine pairs (12). Its functional role is unknown. The helix-turn-helix domain is enigmatic in its own right as the motif canonically mediates protein-nucleic acid binding, which is not expected for PM V in the ER lumen. The structure additionally revealed an unpaired cysteine at the flap over the active site, raising the possibility of redox regulation. Lastly, not visible in the *P. vivax* PM V structure is a predicted flexible loop projecting away from the active site. In *P. falciparum* PM V the corresponding sequence represents a substantially larger and highly charged insert, the role of which we were curious to determine.

Here we utilized our previously described lethal depletion of PM V (9) to probe the above features. Using a genetic rescue method, we found the helix-turn-helix domain, unpaired cysteine, and *P. falciparum*-specific charged insert to be dispensable for parasite growth. In contrast, PM V without a nepenthesin insert was unable to rescue PM V depletion. Further mutagenesis of the nepenthesin insert revealed that the amino acid sequence of the insert as well as some elements of its structure are required for its function. We deployed molecular dynamics simulations to understand how nepenthesin insert mutagenesis impacts the conformational dynamics of PM V. Our simulations show that the nepenthesin insert allosterically modulates positioning of the active site residues and mutation leads to increased prevalence of catalytically nonfunctional states. Taken together, these data broaden our model of how PM V enacts its essential function(s) and provide new insight into the role of the nepenthesin insert in protease biochemistry.

## Results

### PfPM V has several unusual features

We used Phyre2 to model PM V from the genetically tractable *P. falciparum*, using the recently published *P. vivax* PM V structure (Fig. 1) (11, 13). The model closely resembles the *P. vivax* structure (Fig. S1), suggesting that the *P. falciparum* enzyme bears the same unusual motifs found in the *P. vivax* enzyme. The nepenthesin insert is present in PM V across the *Plasmodium* genus, yet is absent in other plasmepsins (Fig. 1A; Fig. S2). The spacing between the four key cysteines that comprise the insert’s structure is conserved among PM V sequences, while the intervening sequences contain some variation (Fig. 1A, Fig. S2). The unpaired cysteine over the active site is completely conserved in PM V sequences across the genus (Fig. 1B). The charged insertion projecting away from the active site is substantially expanded in *P. falciparum* relative to *P. vivax* and includes an unusually dense run of charged amino acids (Fig. 1C). Lastly, the helix-turn-helix domain is conserved throughout the genus, but absent in other *P. falciparum* plasmepsins (Fig. 1D, Fig. S3).

**Figure 1.**
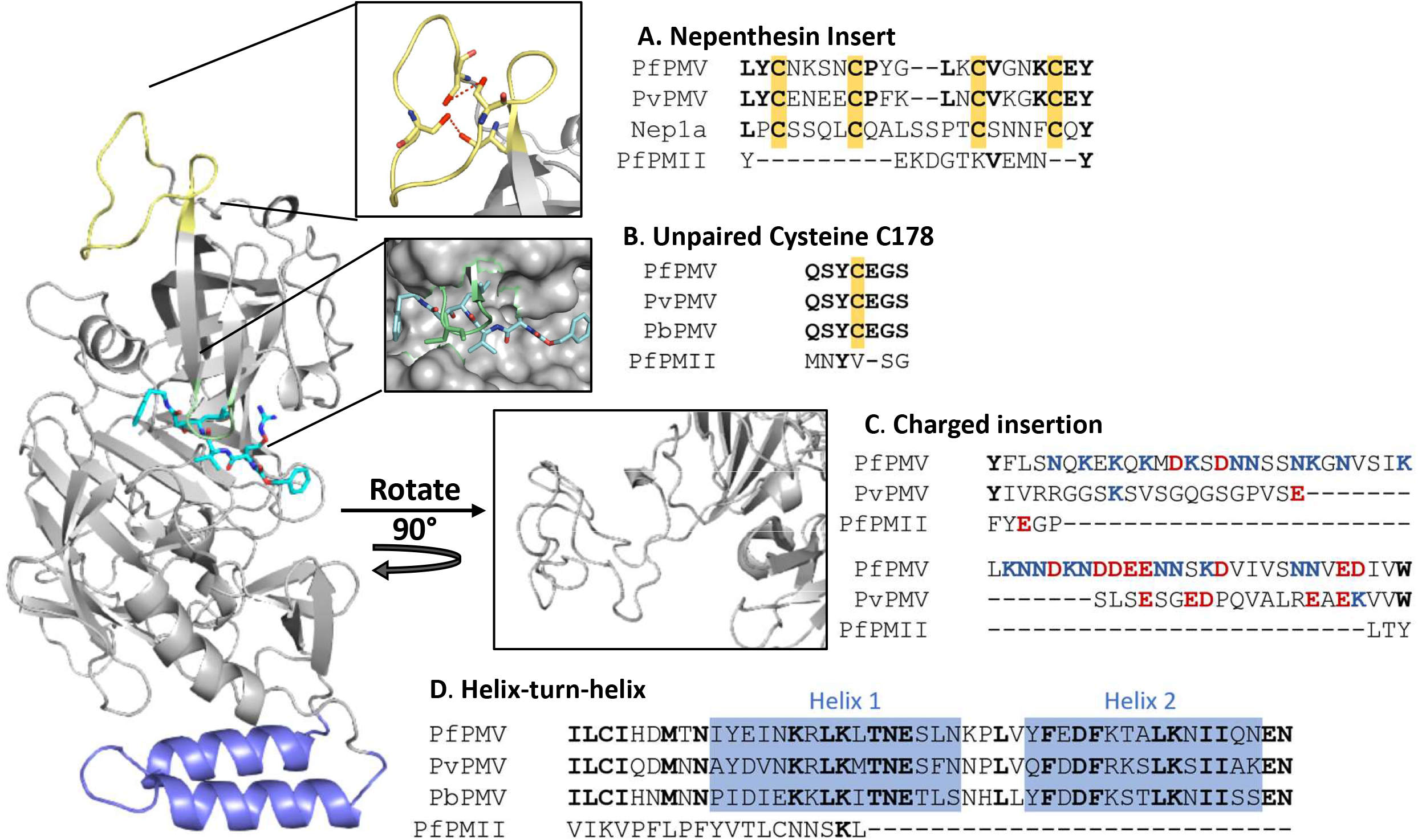
PfPM V structure bears unusual motifs: The structure of PfPM V was modeled using Phyre2 and the published structure from Hodder, Sleebs, Czabotar, et al (2015). The peptidomimetic inhibitor WEHI-842 is shown in bright blue. In this study we investigated (A) the nepenthesin insert, shown in yellow (inset, showing the two Cys-Cys bonds in red), (B) an unpaired cysteine (green) near the active site, (C) a charged insert projecting away from the active site, and (D) a helix-turn-helix domain in blue. At right, the PfPM V sequence aligned next to that of the human parasite *P. vivax*, the rodent parasite *P. berghei*, the paralog plasmepsin II (PfPMII), and where relevant the pitcher plant protease nepenthesin 1a (Nep1a). Shared amino acids are bolded. In (A) and (B) cysteines are highlighted in yellow. In (C) positively and negatively charged amino acids are shown in red and blue respectively.

### Regulatable system for PM V depletion and rescue

To probe the function of these structural motifs we utilized a previously described post-transcriptional depletion of PM V (9). Briefly, we used CRISPR/Cas9 editing to replace the endogenous PM V sequence with a recodonized copy flanked by an N-terminal aptamer and a 10x array of C-terminal aptamers (Fig. 2A). In the mRNA, the aptamers fold into hairpins evolved to bind the Tet Repressor (TetR). TetR is fused to the RNA helicase DOZI, and the binding of the TetR-DOZI fusion protein to the mRNA results in sequestration of the mRNA and depletion of protein levels. The system is controlled by the presence of anhydrotetracycline (aTc). In the presence of aTc, TetR-DOZI preferentially binds aTc and translation continues unabated (14). In the absence of aTc, TetR-DOZI binds the aptamers and translation is suppressed. Knockdown depletes PM V levels greater than 50–fold and is lethal to parasites (9). We attempted to complement the depletion with a second copy of PM V, tagged with C-terminal GFP, and integrated into the neutral *cg6* locus of NF54^attB^ parasites using the previously described attP/attB system (Fig. 2A) (15).

**Figure 2.**
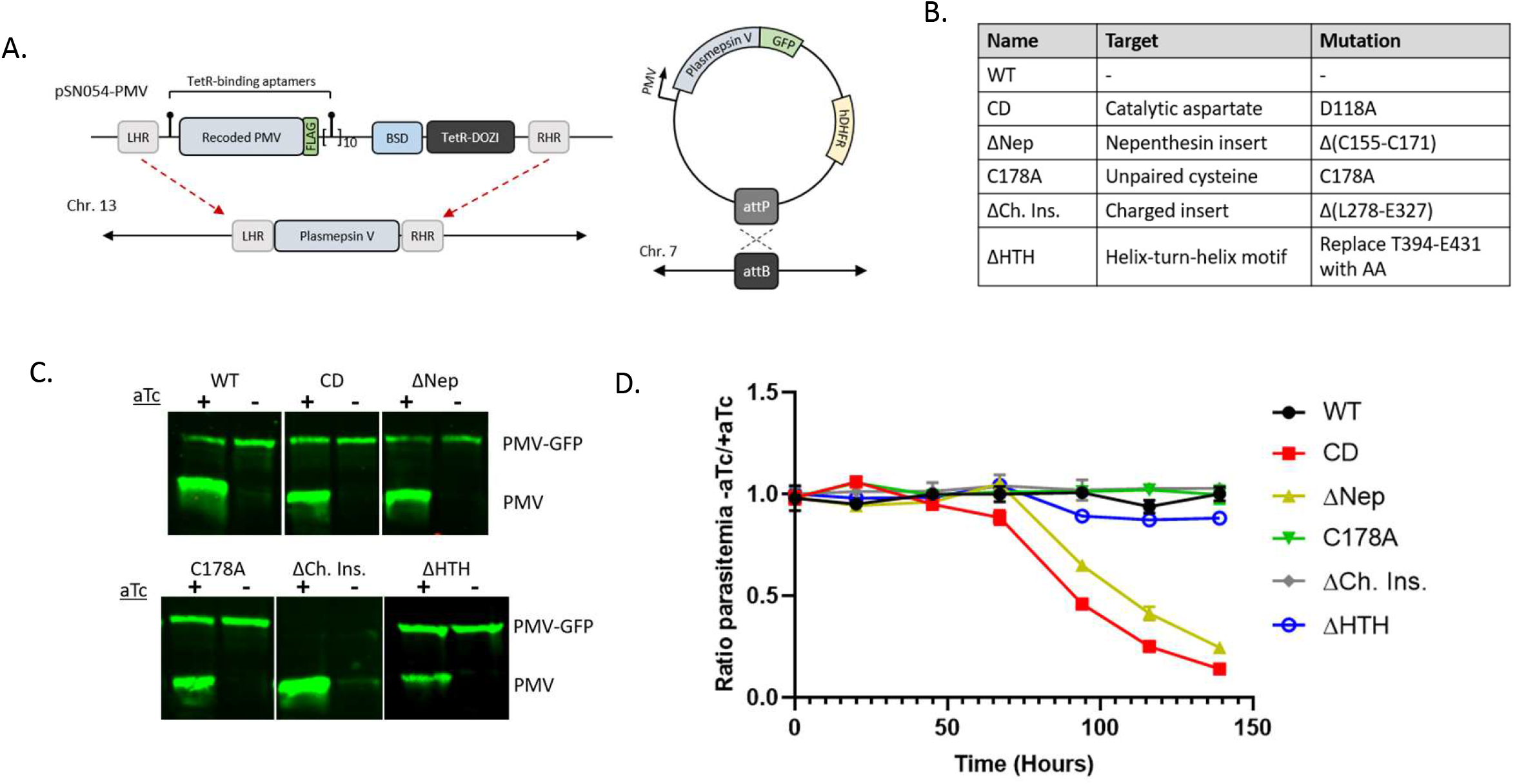
Regulatable system for PM V depletion and rescue: (A) Diagrams of the plasmid system combining pSN054-PMV for tetracyline-regulatable PM V expression, and a GFP-tagged second copy integrated at a genomic attB site and driven by the endogenous promoter. (B) A table of the mutant names in panel C, with the mutations they represent at right. (C) Western blot using anti-PMV antibody. The endogenous copy of PM V is regulatable with aTc; the second copy of mutant PMV-GFP is not. (D) aTc was washed out from asynchronous parasites and growth followed daily by flow cytometry. The experiment was performed three times, each time in technical triplicate. Points represent the mean of technical triplicates from a representative experiment, error bars the standard deviation. Growth curves adjusted for subculturing are shown in Supp. Fig. 5.

To probe the role of PM V structural motifs, we made six PM V-GFP versions: a full-length second copy (WT), a D118A mutant that alters one of the catalytic aspartates ablating catalytic activity (“catalytic dead”, or CD) (11), a complete deletion of the nepenthesin insert from the first to fourth cysteine (ΔNep), replacement of the unpaired flap cysteine with alanine (C178A), deletion of the charged insert (ΔCh. Ins.), and replacement of the helix-turn-helix motif with two alanines (ΔHTH) (Fig. 2B). Expression of each is driven by the PM V promoter, with the exception of ΔHTH, for which cloning issues forced us to use an alternative plasmid driven by the Hsp86 promoter. Parasite lines were selected in the presence of aTc, using blasticidin S (for the knockdown construct) and WR-99210 (for the second copies), and cloned by limiting dilution. The resulting lines each expressed the endogenous PM V as well as the second copy at approximately 2:1 molar ratio (Fig. S4, Fig. 2C); the ΔHTH line, being driven by a stronger promoter, expressed the second copy at an approximately equimolar ratio to the endogenous enzyme (Fig. S4, Fig. 2C). Deletion of the charged insert resulted in an enzyme that was no longer recognized by our previously described PM V monoclonal antibody (16, 17), but was still recognized by anti-GFP (Fig. S4), suggesting that this region contains the binding site for the PM V monoclonal antibody.

### The nepenthesin insert is required for parasite growth

To probe the role of each of these motifs in PM V function, we depleted endogenous PM V by washing out aTc from parasite culture and followed parasite growth daily by flow cytometry. In each case, aTc washout caused substantial depletion of endogenous PM V levels (Fig. 2C). The second copy of WT PM V rescued parasite growth, while CD PM V did not (Fig. 2D). To our surprise, parasite lines expressing PM V lacking the unpaired cysteine, charged insert, or helix-turn-helix domain all grew indistinguishably +/- aTc, suggesting each of these motifs is unnecessary for PM V’s essential function in parasite growth. Only the ΔNep mutant failed to rescue growth in the absence of aTc, suggesting that the nepenthesin insert is required for proper PM V function *in vivo*.

To further dissect the functional contribution of the nepenthesin insert, we mutagenized the insert, attempting to separate its structure from the sequence of its loops. To this end we made three additional mutants: replacing either the first or second cysteine pairs with serines (Nep^C1,3S^ and Nep^C2,4S^ respectively), or replacing the entire nepenthesin insert with the orthologous sequence from the pitcher plant protease Nep1A (Nep^Nep1A^) (Fig. 3A, B), to preserve the insert’s fold while largely altering the lobe sequences. As above, we transfected these as second copies into parasite lines with regulatable endogenous PM V. Washing out aTc from the cultures depleted endogenous PM V levels but had no effect on levels of the second copy (Fig. 3C). After washout, we followed growth daily by flow cytometry (Fig. 3D). PM V with the Nep1A sequence failed to rescue growth, suggesting that the general structure of the insert is not sufficient for its role in PM V function. Of the two cysteine pair mutants, replacement of the first pair had no effect on growth, while replacement of the second pair resulted in a partial growth phenotype – these parasites grew at approximately half the rate without aTc as with. We have previously shown expression of PM V at or above 3% of wildtype levels is adequate to achieve normal growth (9). Therefore the fact that the Nep^C2,4S^ mutant has impaired growth suggests a substantial defect in enzyme function.

**Figure 3.**
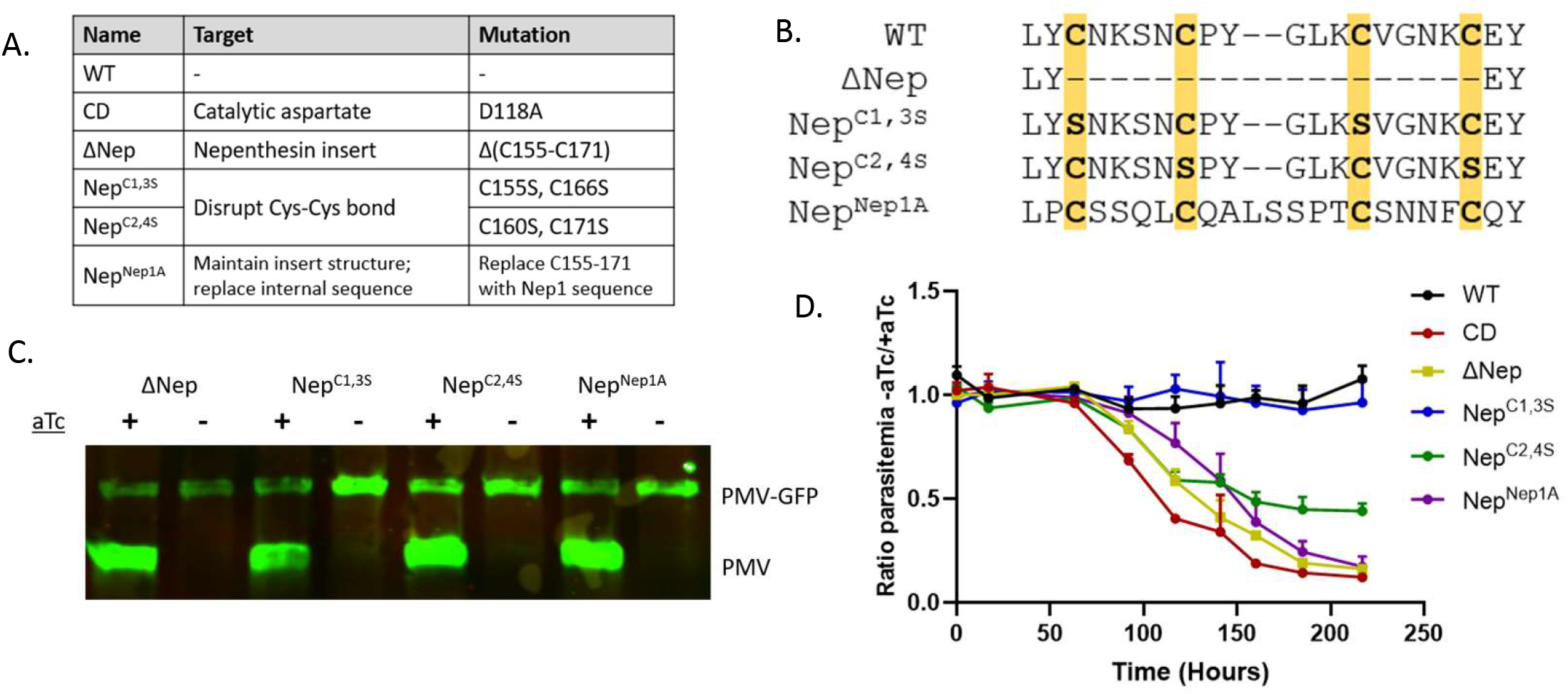
The nepenthesin insert scaffold is not sufficient to rescue nepenthesin loop function. (A) Table of mutants for Figure 3 (B) Sequence of the nepenthesin insert for the mutants described in (A). (C) Western blot with anti-PMV showing regulation of the endogenous gene but not the mutant second copy. (D) aTc was washed out from asynchronous parasites and growth followed daily by flow cytometry. The experiment was performed three times, each time in technical triplicate. Points represent the mean of technical triplicates, error bars the standard deviation. Growth curves adjusted for subculturing are shown in Supp. Fig. 5.

### The nepenthesin insert is required for PM V catalysis

*We* next queried whether the nepenthesin mutants failed to rescue growth due to an effect on PM V catalytic activity, or whether they interfered with some other essential PM V role (e.g. a protein-protein interaction). Recombinant expression in mammalian cells afforded production of some, but not all of the enzyme mutants (Fig. S6). To employ a system where we could compare the full range of mutants, we immunoprecipitated PM V enzyme mutants using anti-GFP from the parasites described above. The eluate contains each mutant PM V without any detectable signal from the endogenous enzyme (Fig. 4A). We then assessed the activity of each mutant enzyme using a previously described FRET peptide assay to assess their activity (6). Briefly, we incubated each enzyme prep with either a “PEXEL” peptide derived from the recognition sequence of the PM V substrate HRP2, or a “mutant” peptide that has a single amino acid change, abrogating PM V recognition. Cleavage of the peptide substrate liberates the fluorophore EDANS from its quencher DABCYL, resulting in fluorescence (Fig. 4B). Incubation with the FRET peptides indicated that the enzymes that could rescue growth (WT, Nep^C1,3S^) were active against PEXEL peptide, but not the mutant peptide (Fig. 4C). The enzymes that could not rescue growth (CD, ΔNep, Nep^Nep1A^) could cleave neither peptide. Strangely, the Nep^C2,4S^ enzyme which partially rescues growth appears inactive in our enzyme assay, perhaps due to the *in vitro* activity buffer not fully recapitulating the environment of the parasite ER, or due to assay insensitivity.

**Figure 4.**
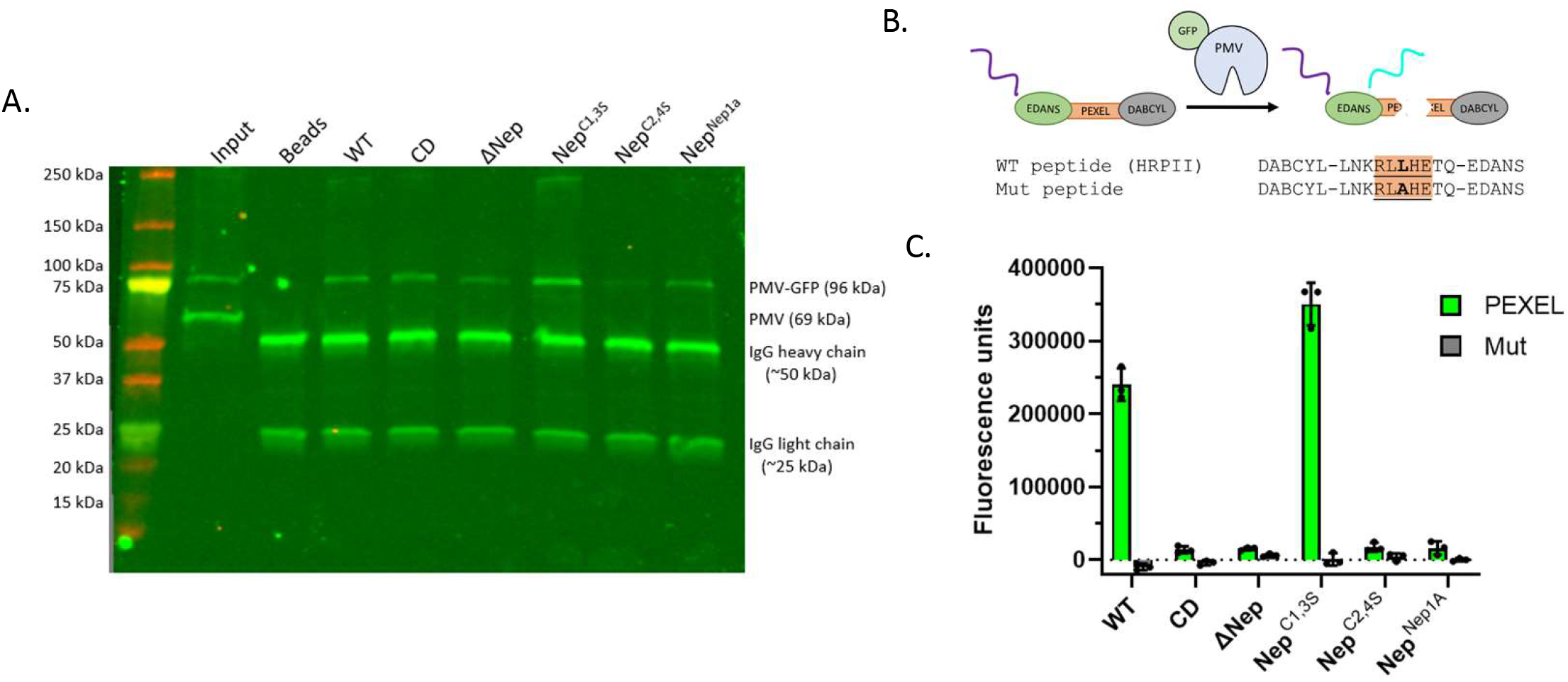
Nepenthesin mutants reduce catalytic activity. (A) Western blot with anti-PM V showing immunoprecipitations of mutant PM V-GFPs. (B) Activity assay scheme. (C) Mutant enzymes were incubated with a PEXEL FRET peptide or a nonrecognized mutant peptide for 12 hours, then fluorescence intensity read as a measure of catalytic activity. Each sample was assayed in technical triplicate, mean fluorescence is shown, with error bars representing standard deviation of the measurements. The experiment was performed three times; a representative experiment is shown.

**Figure 5.**
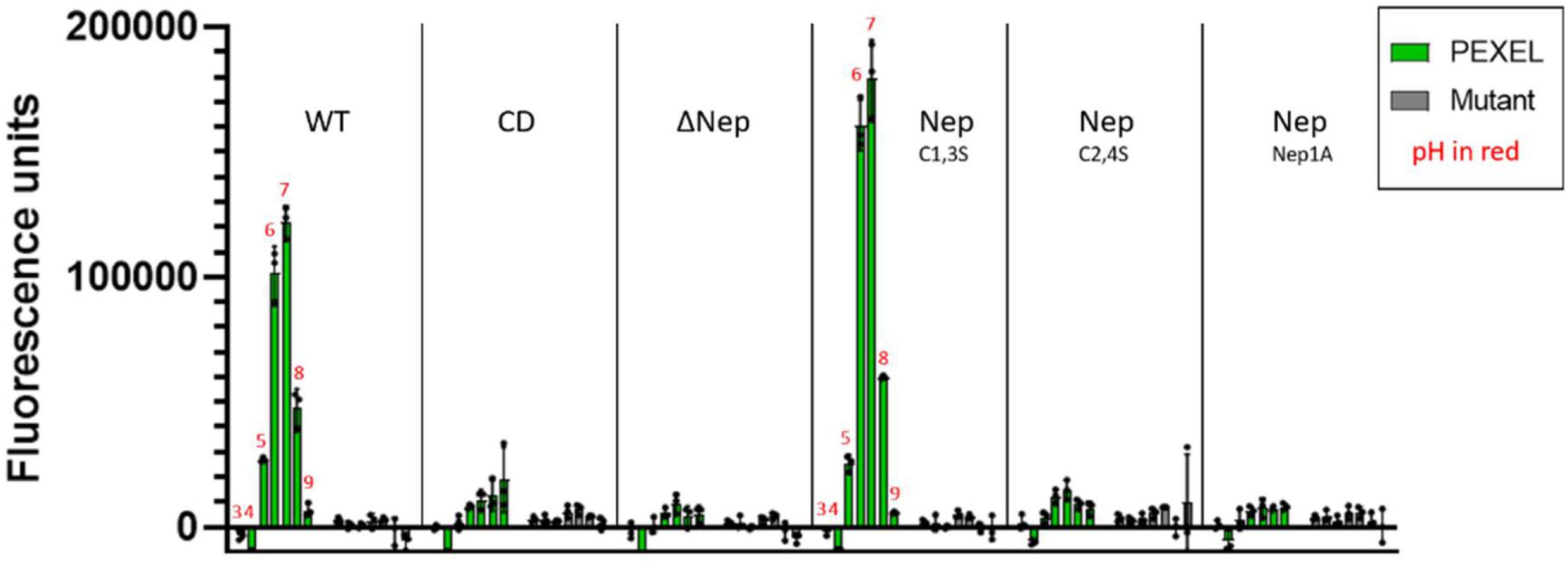
Nepenthesin mutants do not influence pH profile. Activity of each enzyme was assayed as above (Fig. 4C) at buffers that differed only in pH, with pH ranging from 3 to 9 (shown above bars with red numbers). Each sample was incubated with either the PEXEL peptide (green) or the mutant (grey). Each sample was run in technical triplicate. The bar height represents the mean of those three measurements; error bars represent the standard deviation. The experiment was performed twice with similar results, one is shown.

Given the canonical role of pH in activating pepsin-like proteases, and the example of the Nep1A protease being active at low pHs (maximal activity at pH 2-3) (12), we sought to determine whether nepenthesin mutants might have activity at alternative pH ranges. To this end, we probed for PM V activity in buffers ranging from pH 3 to pH 7. We found that the two enzymes active at pH 6.5 (WT and Nep^C1,3S^) were the only ones to display activity at any pH tested, and each displayed a similar pH to profile to that previously described (Fig. 6) (6). Thus, perturbations to the nepenthesin insert do not influence the enzyme’s pH/activity relationship.

**Figure 6.**
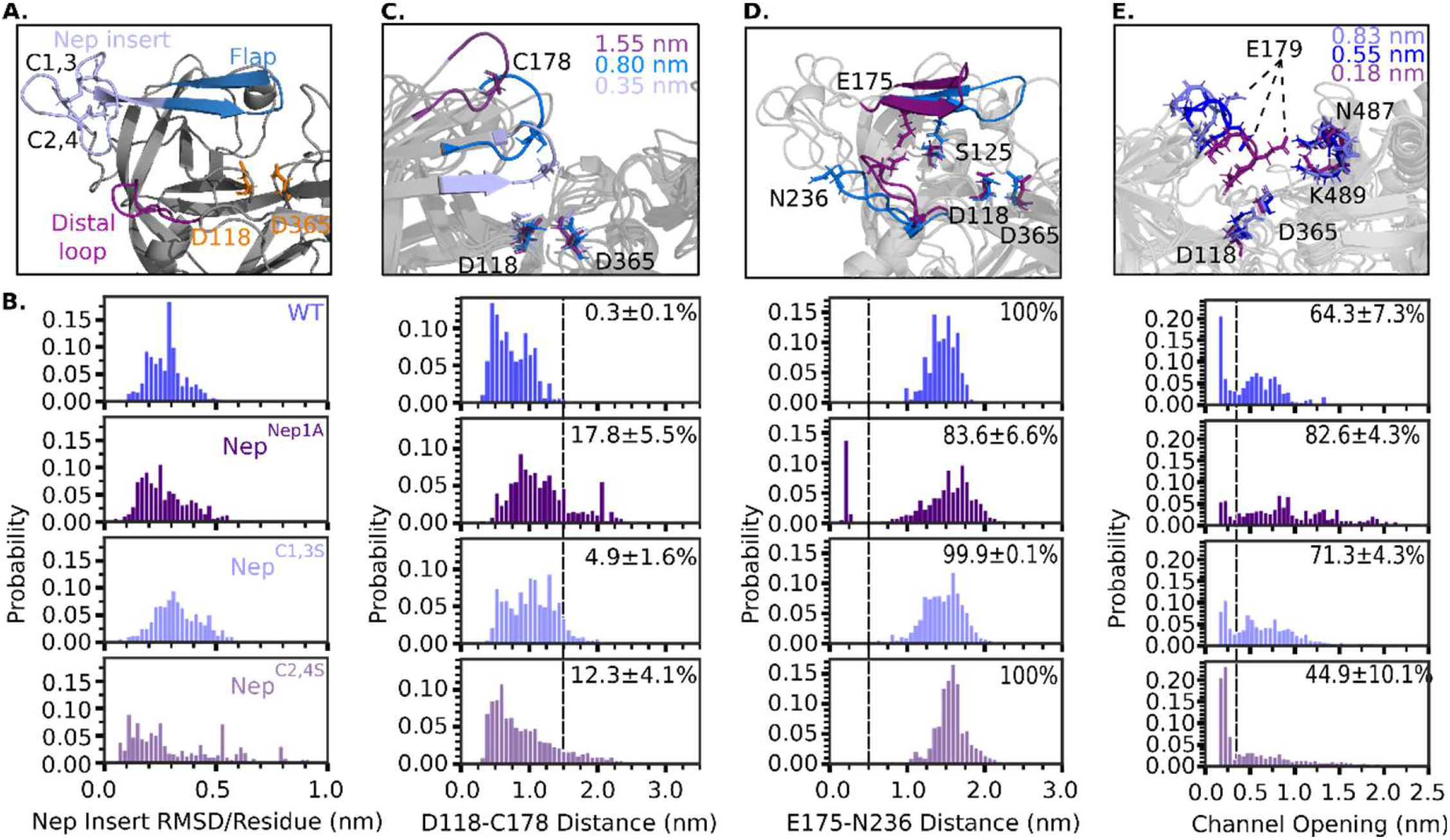
Computational modeling of *P. falciparum* PM V mutants suggests mechanisms for loss of activity: (A) Structural model of *P. falciparum* PM V adapted from PDB: 4ZL4. (B) Impact of Nep insert mutations on flexibility of the Nep insert. Shown is the equilibrium distribution of root mean square deviation (RMSD) distances (in nanometers) for the Nep insert (L153-Y173), normalized per residue. (C) Impact of Nep insert mutations on the opening of the flap covering the active site. Shown are representative structures for different D118-C178 distances for wild-type PM V, as well as the equilibrium distance distributions for each mutant. Quantification is the % of states that occupy a distance larger than a flap opening cutoff of 1.5 nm +/- SD. (D) Impact of Nep insert mutations on distance between E175 (flap) and N236 (distal loop). Shown are representative structures for different channel widths and the equilibrium distributions for each PM V mutant. Quantification is the % of states that occupy a E175-N236 distance > 0.5 nm +/- SD. (E) Impact of Nep insert mutations on active site channel opening distance. Channel width defined as the minimum distance between any atom on side chains 176-181 and 485-490. Quantification is the % of states with their channel more open than 0.4 nm +/- SD. For all quantification, the displayed value is the mean +/- SD as determined by bootstrapping the MSM model by iteratively removing 1 of the 7 trajectories for each mutant.

### The nepenthesin insert allosterically controls PM V flap movement

The C2,4 cysteine pair tethers the nepenthesin insert directly to the N-terminal strand of the flap over the active site (Fig 6A). This tethering has been postulated as a mechanism for functional regulation of PM V and nepenthesin 1, both through protein-protein interactions and protein allostery (11, 12). We posited that mutagenesis of the nepenthesin insert impacts the conformational preferences of the nepenthesin insert leading to the lack of activity. To test this hypothesis, we performed molecular dynamics simulations of WT and Nep^Nep1A^ PM V. We generated initial structures of both *P. falciparum* PM V and Nep^Nep1A^ using the previously solved structure of *P. vivax* PM V (PDB: 4ZL4) as a template and Swiss-Model (18, 19). For both constructs, we ran seven independent unbiased molecular dynamics simulations for 1 μs simulation time per replicate for an aggregate simulation time of 7 μs per mutant. To quantify our results and highlight conformational heterogeneity, we built Markov State Models (MSMs), a statistical framework to analyze molecular dynamics simulations that provides atomistic insights into ensembles of conformations that proteins can adopt (20).

We first investigated how modifications affect the conformational preferences of the nepenthesin insert. We calculated the root mean square deviation per residue of the nepenthesin insert (Fig 6B) and find minimal perturbations to the conformations the insert can adopt, suggesting that the loss of activity in Nep^Nep1A^ is not a result of gross changes in nepenthesin insert conformational preferences.

Curious if mutations to the nepenthesin insert result in allosteric changes that ablate PM V activity, we next turned our attention to processes likely to disrupt catalysis. During PEXEL cleavage, PM V must adopt a flap conformation more open than that seen in the crystal structure to accommodate PEXEL binding. Once PEXEL is bound, we expect the flap to close, trapping the PEXEL motif next to the catalytic aspartates for hydrolysis to occur. We reasoned that if the equilibrium distribution of flap conformations was disrupted in Nep^Nep1A^, this may lead to a loss of catalytic activity. Accordingly, we quantified the extent of flap opening by measuring the minimum distance between D118, one of the active site aspartates, and C178, the unpaired cysteine on the tip of the flap (Fig 6C). In WT PM V, the flap exists in two primary populations, one more closed in a pose similar to that seen in the crystal structure of *P. vivax* PM V, and another slightly expanded. While the Nep^Nep1A^ flap explores these distances, it also explores conformations where the flap has significantly dissociated from the active site. We calculated the probability of the loop exploring a significantly dissociated state (D118-C178 distance > 1.5 nm) and find that WT has an equilibrium probability of 0.3 ± 0.1% for these states, whereas Nep^Nep1A^ has a probability of 17.8 ± 5.5%.

While characterizing the motion of the flap, we noticed extensive hydrogen bonding between the flap and the distal cavity of the active site. Previous studies have highlighted that loss of H-bonding between the flap and active site adjacent pockets of aspartic proteases dramatically impacts the catalytic activity of aspartic proteases (21). Notably, in WT PM V, S125 spends the majority of our simulations in a hydrogen bond with either E175 or H212 (mimicking the H-bond network between Y77 and W41 in PM II) (22, 23). In Nep^Nep1A^, H-bonding to S125 is significantly disrupted (Fig 7). In Nep^Nep1A^, when S125 is not in a H-bond the flap is more likely to explore conformations where the flap is skewed significantly to the side. This skewing allows the stable formation of a H-bond between E175 on the flap and N236 in the distal loop (Fig 6A,C), a configuration not observed in our WT simulations.

**Figure 7:**
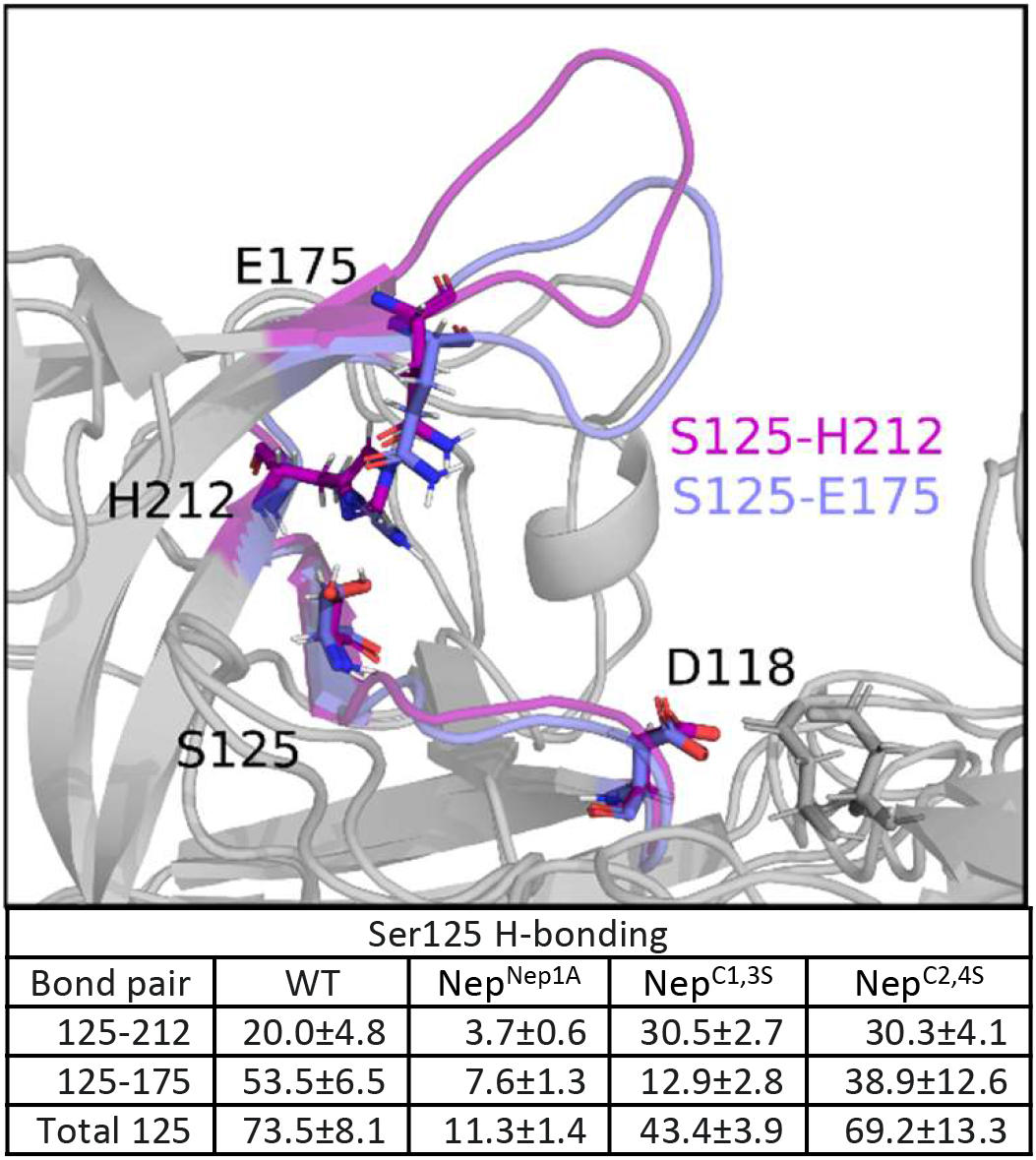
H-bonding network of PM V. Displayed are the most probable H-bonds for WT-PM V. Quantification is the probability of a hydrogen bond existing according to the Wernet Nilsson criteria (21). Displayed is the mean +/- standard deviation from a MSM bootstrapped by iteratively leaving one simulation replicate out.

We also wanted to understand why Nep^C1,3S^ maintains activity whereas Nep^C2,4S^ results in a loss of activity. In the crystal structure of *P. vivax* PM V, the Nep^C1,3^ disulfide bond holds the nepenthesin insert together, whereas Nep^C2,4^ anchors the nepenthesin insert to the active site flap. As before, we simulated 7 independent replicates for 1 μs each for Nep^C1,3S^ and Nep^C2,4S^ and built a MSM to quantify our results. While the Nep^C1,3S^ nepenthesin insert had similar dynamics to Nep^Nep1A^, Nep^C2,4S^ was much more flexible than all other mutants (Fig 6B). Similarly, the Nep^C1,3S^ flap explores conformations similar to WT, whereas Nep^C2,4S^ explores conformations more similar to Nep^Nep1A^ (Fig 6C). We expected that the increased flexibility of the nepenthesin insert resulting from Nep^C2,4S^ would allow the flap to similarly interact with the distal loop whereas Nep^C1,3S^ would, like WT PM V, never interact with the distal loop. Surprisingly, neither Nep^C1,3S^ nor Nep^C2,4S^ interact with the distal loop (Fig 6D).

Having explored two mechanisms of catalytic prevention – active site closure by flap clamping/dissociation and loss of substrate interaction through flap pinning – we wanted to check the final mechanism by which the active site may be occluded: through closure of the active site. Accordingly, we measured the minimum distance between the active site flap and the loop opposite the active site which forms the active site channel (Fig 6E). In our simulations of WT PM V, the channel is “closed” (defined as roughly the width of a peptide backbone, minimum channel distance < 4Å) 35.7 +/- 7.3% of the time. Nep^Nep1A^ and Nep^C1,3S^ have closed channels 17.4 +/- 4.3% and 28.7+/- 4.3% of the time, whereas Nep^C2,4S^ has a stark increase in the population of closed channel conformations, occurring 55.1 +/-10.1% of the time.

## Discussion

Here we used parasite genetics and molecular dynamics simulations to investigate the relationship between PM V’s unusual structure and its function in parasite biology. We find that the PMV nepenthesin insert is required for parasite survival and for catalysis of a substrate peptide *in vitro*, whereas a helix-turn-helix motif, unpaired cysteine near the active site, and large charged insert were dispensable for parasite survival (and presumably for PM V catalysis).

The most intriguing motif in our story is the nepenthesin insert. This motif was originally described by Athauda, et al. (12) as a 22-residue insertion unique to nepenthesin-like aspartic proteases (and absent from their closest plant orthologues). The finding that PM V bears a similar insert was a surprise (17), given that it has little in common with the nepenthesin-type aspartic proteases. The nepenthesin proteases are digestive enzymes, secreted into the pitcher plant lumen, and activated at very acidic pH (optimal activity at pH ^~^2.5) to participate in the digestion of insect prey material. Both nepenthesin I and nepenthesin II have broad substrate specificity, and can cleave before or after many residues (24). In contrast PM V is retained in the ER at much higher pH (optimal activity at pH ^~^6.5) and has only been described to cleave C-terminal to RxL (6, 25). The same motif in both sets of aspartic proteases, but not in others, defies obvious explanation.

Athauda, et al. speculated that the nepenthesin insert may influence localization or regulation of the enzyme; the subsequent finding of a nepenthesin insert in PM V makes at least the former seem less likely. Hodder, Sleebs, Czabotar, et al. suggest that the insert may be involved in interaction with other proteins, perhaps the PEXEL substrates themselves (11). We have not tested this hypothesis here, but can comment that the *Toxoplasma gondii* ortholog Asp5 cleaves an RRL motif that is very similar to the RxL recognized by PM V (18–20) yet appears to have a large insert between the third and fourth cysteine of its orthologous nepenthesin insert sequence, giving spacing of CX_5_CX_3_CX_19_C compared to the CX_4_CX_5_CX_4_C spacing in *Plasmodium*. One could imagine that this extra 15-amino acid run would disrupt the nepenthesin insert, however without structural information on Asp5, this remains unclear. We have speculated that the insert may serve as a docking point for chaperones (26), yet to our surprise all nepenthesin insert mutations that disrupted parasite growth also blocked catalysis *in vitro*, suggesting that the insert contributes to substrate recognition, access, or cleavage. Our molecular dynamics simulations suggest that mutation of the nepenthesin insert results in a loss of catalytic activity for three reasons. First, Nep^Nep1A^ and Nep^C2,4S^ increase the likelihood that the active site flap has dissociated from the active site and is therefore unable to hold a substrate in the active site to enable hydrolysis. Second, Nep^Nep1A^ results in an extra flexible flap that becomes stably pinned to the distal loop via H-bonding. This pinning would likewise preclude the flap holding a substrate in the active site. Finally, Nep^C2,4S^ results in an increased probability of a closed active site channel, which would prevent substrate binding.

As for the other structural motifs studied here, the helix-turn-helix, unpaired cysteine, and large charged insertion each appear dispensable for parasite growth in culture. Helix-turn-helix domains canonically mediate protein-DNA interactions, but have also been described to have roles in protein-RNA interactions, protein-protein interactions, and structural support for enzymatic domains (27). In this case, the conservation of the domain in PM V sequences throughout the genus suggests a significant role for this motif. Our data are limited to blood-stage parasites in culture; perhaps studies in different *Plasmodium* life stages or animal models of infection would reveal a yet unappreciated role.

Similarly, the unpaired cysteine appears dispensable for parasite growth. The enzyme domain of PM V is rich in cysteines, with 15 cysteines arranged into seven pairs – compared to plasmepsin II which has five cysteines arranged into two pairs (11, 28). This disulfide-rich architecture has earned PM V classification into MEROPS peptidase subfamily A1B, with type member Nep1, rather than the canonical pepsin subfamily A1A (which includes all other plasmepsins) (29). In A1B subfamily members, the richer network of disulfide bonds has been hypothesized to bolster the enzyme structure against harsh conditions (12). This makes sense for the nepenthesins, which must act in acidic digestive conditions; perhaps for PM V this enhanced structure protects against the oxidizing environment of the ER. The unpaired cysteine C178 has been hypothesized to have a role in metal binding, and indeed recombinant PM V with the same C178A mutation we describe here was shown to have altered susceptibility to inhibition by Hg^2+^ ions (30). The physiological implications of this observation remain unresolved, as we see no impact of this mutation in parasite culture. C178 and its surrounding amino acids are perfectly conserved across the *Plasmodium* genus. Whether that conservation presages a functional contribution at another part of the parasite life cycle, we cannot yet say.

Lastly, the *P. falciparum* genome is inexplicably full of insertions of charged and presumably poorly structured sequences in proteins that lack these insertions in the rest of the genus (31, 32). These insertions often form surface loops with no known function (32). Muralidharan et al. showed that deletion of an asparagine-rich repeat from the proteasome subunit rpn6 had no discernible effect on the protein’s essential function (33), and suggested that the *P. falciparum* Hsp110 may be so effective at preventing aggregation of repetitive prionogenic sequence as to nearly eliminate the fitness cost of maintaining these sequences (34). Here, we describe a similar example: a surface-exposed, low complexity loop, rich in charged amino acids, that has no apparent role in PM V function. Whether these sequences play roles in certain setting or are merely an evolutionary quirk merits further exploration.

In summary, our data provide insight into the essential function of PM V in the *P. falciparum* intraerythrocytic cycle. Given the unusual nature of this enzyme’s structure, this also provides atomistic insights into the contribution of the nepenthesin insert and its role in side-chain heterogeneity that modulates conformational dynamics of PM V, which we hope will support further investigation into this motif and A1B subfamily proteases more broadly. Future studies will also highlight role of side-chain heterogeneity in flap opening and designing novel ‘open flap’ inhibitors targeting PM V (35, 36).

## Acknowledgements

We thank members of the Goldberg lab for invaluable suggestions over the course of this project, particularly Eva Istvan and Sumit Mukherjee for sharing their expertise in protein biochemistry, and Barbara Vaupel for assistance making plasmids.

## Materials & Methods

### Modeling and Alignments

The *P. falciparum* PM V structure was modelled using Phyre2. Mutant sites were selected by aligning the *P. falciparum* PM V structural model with the published *P. vivax* PM V (11) one using PyMOL (v 2.3). Sequence alignments were performed using reference genome sequences retrieved from PlasmoDB (37) and aligned with Clustal Omega (38). Structures for molecular dynamics simulations were generated using SWISS-MODEL (18, 19) using the published structure of *P. vivax* PM V (11) as a homology model.

### DNA constructs

To modify the parasite genome, we used the previously described AttP/AttB recombinase system which utilizes a recombinase-expressing plasmid pINT (15) along with a donor plasmid that gets integrated into the genome at the *cg6* locus. Here, our donor plasmid was a derivative of pEOE (34, 39) modified to include the attP sequence using the QuikChange Lightning Multi Site Directed Mutagenesis kit and the primer 1 (see Sup. Table 1 for list of primers used in this study) as in (40) to create the plasmid pEOE-AttP. To make PM V complementation vectors, the full PM V sequence and its 5’ UTR were PCR amplified from the NF54 genome using primers **2** and **3,** gel purified, and inserted by Infusion (Clontech) into pEOE-AttP cut with AatII and AvrII. The resulting plasmid, pEOE-AttP-PMV_prom_-PMV-GFP, was then mutagenized using the QuikChange Lightning Multi Site Directed Mutagenesis kit and primers **5, 6, 7, 8, 10, 11,** or **12** to make rescue plasmids with each PM V mutant in Fig. 2B and Fig. 4A. The exception, as noted above, was the ΔHTH mutant. This plasmid was instead made by amplifying the plasmepsin V sequence with primers **2** and **4,** and inserting into pEOE-AttP cut with XhoI and AvrII (maintaining the existing *hsp86* promoter), then mutagenizing as above but with primer 9. Plasmids sequences were verified by Sanger sequencing (Genewiz; South Plainfield, NJ) using primers **13, 14, 15,** and **16.**

For mammalian cell expression, we ordered a synthesized sequence for the soluble domain of PM V (V36-S520) codon optimized for expression in human cells (Integrated DNA Technologies). This sequence was PCR amplified with primers **17** and **18,** then inserted into KpnI–digested pHLsec (41) by Infusion. Plasmids for expressing mutant enzyme were made as above, using primers **19, 20, 21, 22,** and **23.** Completed plasmids were verified by Sanger sequencing using primers **24** and **25.**

### Parasite lines and culture

*P. falciparum* strain NF54^attB^ (15) was cultured in RPMI 1640 (Gibco) supplemented with 0.25% (w/v) Albumax (Gibco), 15 mg/L hypoxanthine, 110 mg/L sodium pyruvate, 1.19 g/L HEPES, 2.52 g/L sodium bicarbonate, 2 g/L glucose, and 10 mg/L gentamicin as well as human red blood cells at 2% hematocrit. We obtain deidentified human red blood cells from St. Louis Children’s Hospital blood bank as well as American Red Cross Blood Services (St. Louis, MO). Cultures were grown in the presence of 100 nM anhydrotetracycline (aTc; Cayman Chemical) unless noted otherwise.

To generate the lines used in this study, we combined 50 μg of each pEOE-AttP-PMV plasmid mentioned above with 50 μg pINT in cytomix (120 mM KCl, 0.15 mM CaCl_2_, 2 mM EGTA, 5 mM MgCl_2_, 10 mM K_2_HPO_4_, 25 mM HEPES) and used that to resuspend a pellet from 5 mL parasite culture with ^~^5% young ring-stage parasites. We then electroporated in a 2mm gap cuvette with a BioRad Gene Pulser II (0.31 kV, 0.950 μF, “High Cap”, “Infinite” resistance on Pulse Controller II). After 24 hours we added 10 nM WR99210 as a selection agent, then monitored cultures thrice weekly by thin smear until parasites emerged. Clonal lines were isolated by limiting dilution and verified by PCR and western blot.

### Western blotting

Western blots were performed as in (9). Briefly, RBCs were lysed in PBS + 0.035% saponin, washed to remove hemoglobin, and the resulting parasite pellets solubilized with RIPA (50 mM Tris pH 7.4, 150 mM NaCl, 0.1% SDS, 1% Triton X-100, 0.5% DOC) plus HALT-Protease Inhibitor Cocktail (Thermo Fisher), cleared by centrifugation to remove hemozoin crystals, and then diluted into 4X Protein Loading Buffer (Licor), and boiled at 99°C. Prepared lysates were separated on Mini-PROTEAN TGX Gels (4-15%, BioRad), transferred to 0.45 μm nitrocellulose, blocked in StartingBlock blocking buffer (ThermoFisher Scientific), and probed with mouse anti-PMV 1:50 (16) and/or rabbit anti-GFP (1:100; Abcam, ab6556), followed by secondary antibodies goat anti-mouse IRDye 800CW 1:10,000 (Licor) and donkey anti-rabbit IRDye 680RD 1:10,000 (Licor). Stained membranes were imaged on a Licor Odyssey platform, and brightness/contrast adjusted for display using ImageStudio Lite 5.2.

### Growth assays

To assess the ability of each mutant enzyme to rescue parasite growth, cultures were washed to remove aTc (3 washes, 5 minutes each) and resuspended in medium containing either 100 nM aTc or an equal volume of DMSO (to deplete endogenous PM V). Growth was monitored daily by flow cytometry using a BD FACS Canto and measuring fluorescence from acridine orange (1.5 μg/mL in PBS). Parasites were subcultured as needed to prevent overgrowth. Parasitemias were quantified using BD FACSDiva, and are displayed as a ratio of the parasitemias of the DMSO culture (“-aTc”) to the aTc culture (+aTc). Graphs were made using Prism (Version 9).

### Peptide assays

PM V activity assays were performed as in (6). Briefly, parasite cultures were pelleted and then RBCs lysed in PBS + 0.035% saponin (10 mL PBS + saponin per 50 mL culture) for 15 min at 4°C. Parasites were pelleted at 15,500 g for 3 min, resuspended in PBS, pelleted again at 15,500 g for 3 min, then frozen at −80°C for days to weeks until the day of the assay. Eventually pellets were thawed and solubilized in PBS + 0.5% Triton X-100, sonicated to disrupt the pellet, then incubated at 4°C for one hour. Lysates were then incubated for one hour at 4°C with Dynabeads-Protein A or Dynabeads-Protein G (ThermoFisher) that had been bound to anti-GFP (clone 3E6, ThermoFisher). Beads were collected on a magnet stand, washed five times in cold PBS + 0.02% Tween-20, and then incubated directly with either the fluorigenic PEXEL peptide DABCYL-LNKRLLHETQ-EDANS, or the mutant peptide DABCYL-LNKRLAHETQ-EDANS at 1 μM in PM V activity buffer (50 mM Tris-Malate, 50 mM NaCl, 0.05% Triton X-100, pH 6.5 unless otherwise noted) for 12 hours at 37°C. Total fluorescence was read on a multimodal plate reader (PerkinElmer Envision). The pH profiling assay was performed as above, but with PM V activity buffer brought to pH 3 with hydrochloric acid, or brought up to pH 4, 5, 6, 7, 8, or 9 with sodium hydroxide.

### Mammalian cell expression

To express PM V in mammalian cells, we transfected HEK293-F cells (ThermoFisher) using a modified version of the protocol in (41). Briefly, HEK293-F cells were cultured in FreeStyle 293 Expression Medium (ThermoFisher). A days before transfection, cells were seeded at 6 x 10^5^ cells/mL, 100 mL/transfection. The next day, with cells ^~^10^6^ cells/mL, transfection was performed by combining 300 μg polyethylenimine (PEI, 25 kDa branched) with 15 mL FreeStyle 293 Expression Medium, slowly adding 100 μg of the plasmid of interest, allowing to sit for 15 min at room temperature, then adding all to the flask of cells. Cells were incubated shaking at 37°C for 48 hours, then supernatants collected and frozen at −80 °C for up to a week.

On the day of purification, supernatants were thawed and bound to a column of nickel agarose beads (GoldBio; St. Louis, MO), washed five times in wash buffer (20 mM imidazole, 20 mM H_2_NaO_4_P*2H_2_O, 500 mM NaCl, pH 7.4), and eluted three times in elution buffer (500 mM imidazole, 20 mM H2NaO4P*2H2O, 500 mM NaCl, pH 7.4). Elutions were then prepared for western blot as above (see “Western blotting”) or used directly for activity assay as above (see “Peptide assays”).

### Molecular Dynamics Simulations

All simulations were prepared using Gromacs 2020 (42). Initial structures were placed in a dodecahedral box extending 1.0 nm beyond the protein in all dimensions. These systems were subsequently explicitly solvated in water (tip3p) along with 0.1M NaCl (43). Systems were energy minimized with a steepest descents algorithm until the maximum force fell below 100 kJ mol^-1^ nm^-1^, using a step size of 0.01nm and a cut-off distance of 1.2 nm for the neighbor list, Coulomb interactions, and van der Waals interactions. All simulations used the AMBER03 force field.

Systems were subsequently run for a total simulation time of 1 μs each using a 4 fs time step using virtual sites. Cutoffs of 1.1 nm were used for the neighbor list, with 0.9 for Coulomb and van der Waals interactions. The Verlet cut-off scheme for neighbor lists was used. Long range interactions were calculated using the particle mesh Ewald method with a Fourier spacing of 0.12 nm. The system temperature was held constant at 300K using the stochastic velocity rescaling (v-rescale) thermostat, and pressure was coupled with Parrinelo-Rahman barrostat (44). 7 independent simulations were run for each mutant: wild-type PM V, Nep^Nep1A^, Nep^C1^,^3S^, and Nep^C2,4S^.

Each PM V mutant was clustered independently using root-mean-square-deviation (RMSD). Clustering was performed using the k-hybrid clustering algorithm to 3000 cluster centers. Markov state models (a network representation of the free-energy landscape (45, 46)) were built using our python package, enspara (https://github.com/bowman-lab/enspara) (45). To estimate the errors associated with our model, we iteratively built MSMs, each time leaving one of our trajectories. Displayed quantifications are the mean of the 7 MSMs and +/- the standard deviation of these MSMs. Transition probability matrices were produced by counting the transitions between each state of the cluster space, adding a prior count of 1/n where n is the total number of states, and row-normalizing as is common (45, 46). Equilibrium populations were calculated as the eigenvector of the transition probability matrix with an eigenvalue of one. All subsequent analysis was performed using the built-in tools of MDTraj (47), and graphs were produced in Matplotlib (48).

