## Supplemental Figures for "A nepenthesin insert allosterically controls catalysis in the malaria parasite protease plasmepsin V"

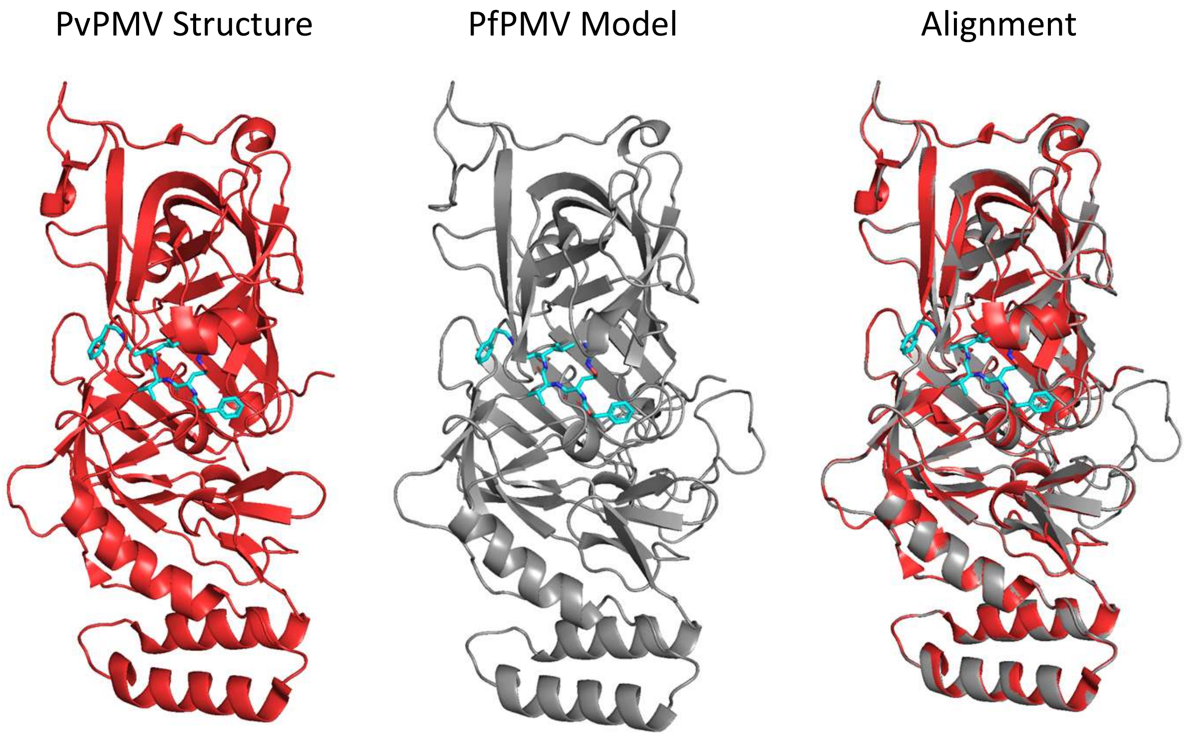


**Supplemental Figure 1 –** Alignment of the experimentally determined *P. vivax* PM V (PvPMV) structure (left, red) and the *P. falciparum* PM V (PfPMV) model predicted by Phyre2 (grey, center). At right, the two structures are aligned, showing their gross structural similarity. Peptidomimetic inhibitor WEHI-842 is shown in blue.

```
PfPMV      YNLNYSKTSSILYCNKSNCPYGLKCVGNKCEYLQSYCEGSQIYGFYFSDIVTLPSYNNK-
PvPMV      FNLNNSKTSSILYCENEECPFKLNCVKGKCEYMQSYCEGSQISGFYFSDVVSVVSYNNE-
PkPMV      FNLNNSKTSSILYCENEKCPYNLNCVNGKCEYLQSYCEGSQISGFYFSDVVTMTSYSNE-
PbPMV      FNLNNSSTSSILYCNDNICPYNLKCVKGRCEYLQSYCEGSRINGFYFSDIVRLESNNNTK
PyPMV      FNLNNSSTSSVLYCNDNTCPYNLKCVKGRCEYLQSYCEGSRINGFYFSDVVKLESTNNTK
PfPMI      FNPNKSRTFTKNLKNNQ-E----------SVYTYIQYGTGTSILEQSYDDVYLK-GLKIK
PfPMII     YKRTKSFVYKYDKKG---L----------PSVIEIFYLSGKIVAFEGYDTIYLGKKLKIP
PfPMIII    YDSSKSKTYEKD-----------------DTPVKLTSKAGTISGIFSKDLVTIG-KLSVP
PfPMIV     YDASASKSYEKD-----------------GTKVEISYGSGTVRGYFSKDVISLG-DLSLP
PfPMVI     FNPNKSRTFTKNLKNNQ-E----------SVYTYIQYGTGTSILEQSYDDVYLK-GLKIK
PfPMVII    YKRTKSFVYKYDKKG---L----------PSVIEIFYLSGKIVAFEGYDTIYLGKKLKIP
PfPMVIII   YDHKISKNYKLVKK---------------KDPVEILFGTGEIHIAYVTDDIHLG-DIKVK
PfPMIX     YNHKLSSSFKYYEP---------------HTNLDIMFGTGIIQGVIGVETFKIG-PFEIK
PfPMX      YDPNKSKTFRRSFI---------------EKNLHIVFGSGSISGSVGTDTFMLG-KHLVR
```

**Supplemental Figure 2 –** Sequence alignment of the area around the nepenthesin insert for PM V from several species, as well as all *P. falciparum* plasmepsins. Alignments were performed using Clustal Omega. Residues shared among the PM V sequences are bolded. The nepenthesin insert is highlighted in yellow.

```
PfPMV   ----------------------------MNNYFLRKENFFILFCFVF------VSIFFVSNVTIIKCNNVENK-----ID   41
PvPMV   --------------------------------------------------------------------------------    0
PbPMV   ------------------------------------------MCFSK--LYNFITYFIIINILV-QA-------------
PfPMI   MALSIKEDFSSAFAKNESAVNSSTFNNNMKTWKIQ-KRFQILYVFFFLLITGALFYYLIDNVLFPKNKKINEIMNTSKHV

PfPMV   NVGKKIENVGKKIGDMENKNDN------------VE-NKNDNVGNKNDNVKNASSDLYKYKLYGDIDEYAYYFLDIDIGK  109
PvPMV   MVGASLGPPGRG--SLSRLIRLVICV--------LTLCALSVQGRSESTEGHSKDLLYKYKLYGDIDEYAYYFLDIDIGT   70
PbPMV   ----------------NTEN--------------------EDILN---KNSEKNEEIYKYKLYGDIDEYAYYFMDINIGT
PfPMI   IIGFSIENSHDRIMKTVKQHRLKNYIKESLKFFKTGLTQKPHLGNAGDS-------V----TLNDV-ANVMYYGEAQIGD

                                                      Nepenthesin Insert      Flap
PfPMV   PSQRISLILDTGSSSLSFPCNGCKDCGIHMEKPYNLNYSKTSSILYCNKSNCPYGLKCVGNKCEYLQSYCEGSQIYGFYF  188
PvPMV   PEQRISLILDTGSSSLSFPCAGCKNCGVHMENPFNLNNSKTSSILYCENEECPFKLNCVKGKCEYMQSYCEGSQISGFYF  150
PbPMV   PGQKLSLIVDTGSSSLSFPCSECKDCGVHMENPFNLNNSSTSSILYCNDNICPYNLKCVKGRCEYLQSYCEGSRINGFYF
PfPMI   NKQKFAFIFDTGSANLWVPSAQCNTIGCKTKNLYDSNKSKTYEK------------DGTKVEM----NY-VSGTVSGFFS

PfPMV   SDIVTLPSYNNK--NKISFEKLMGCHMHEESLFLHQQATGVLGFSLTKP---NGVPTFVDLLFKHTPSL-KPI---YSIC  259
PvPMV   SDVVSVVSYNNE---RVTFRKLMGCHMHEESLFLYQQATGVLGMSLSKP---QGIPTFVNLLFDNAPQL-KQV---FTIC  220
PbPMV   SDIVRLESNNNTKNGNITFKKHMGCHMHEEGLFLHQHATGVLGLSLTKP---KGVPTFIDLLFKSSPKL-NKI---FSLC
PfPMI   KDIVTIANLSFP----YKFIEVTDTNGF-EPAYTLGQFDGIVGLGWKDLSIGSVDPVVVEL--KNQNKIEQAVFTFYLPF

                                 Unstructured region
PfPMV   VSEHGGELIIGGYEPDYFLSNQKEKQKMDKSDNNSSNKGNVSIKLKNNDKNDDEENNSKDVIVSNNVEDIVWQAITRKYY  339
PvPMV   ISENGGELIAGGYDPAYIVRRGGSKSVSGQ------GSGPVSESLS--------ESGEDPQVALREAEKVVWENVTRKYY  286
PbPMV   ISEYGGELILGGYSKDYIVKEVSIDEKKDNIEH--------------------NKNENINSINKSIVDGILWEAITRKYY
PfPMI   DDKHKGYLTIGGIEDRFYEGQLT------------------------------------------------YEKLNHDLY

                                                                     Helix-turn-helix
PfPMV   YYIKIYGLDL-YGTNIM-DKKELDMLVDSGSTFTHIPENIYNQINYYLDILCIHDMTNIYEINKRLKLTNESLNKPLVYF  417
PvPMV   YYIKVRGLDM-FGTNMMSSSKGLEMLVDSGSTFTHIPEDLYNKLNYFFDILCIQDMNNAYDVNKRLKMTNESFNNPLVQF  365
PbPMV   YYIRVKGFQL-FGTTFSHNNKSMEMLVDSGSTFTHLPDDLYNNLNFFFDILCIHNMNNPIDIEKKLKITNETLSNHLLYF
PfPMI   WQVD---LDLHFGNL---TVEKATAIVDSGTSSITAPTEFLNKFFEGLDVVKIPFLP----------LYITTCNNPKLPT

PfPMV   EDFKTALKNIIQNENLCIKIVDGVQCWKSLENLPNLYITLSNNYKMIWKPSSYLYKKESF---WCKG--LE-KQVNNKPI  491
PvPMV   DDFRKSLKSIIAKENMCVKIVDGVQCWKYLEGLPDLFVTLSNNYKMKWQPHSYLYKKESF---WCKG--IE-KQVNNKPI  439
PbPMV   DDFKSTLKNIISSENVCVKIADNVQCWRYLENLPNIYIKLSNNTKLVWQPSSYLYKKESF---WCKG--LE-KQVNDKPI
PfPMI   LEFRSATNVY------------------------------------TLEPEYYLQQIFDFGISLCMVSIIPVDLNKNTFI

PfPMV   LGLTFFKNKQVIFDLQQNQIAFIESKCPSNLTSSRPRTFNEYREKENIFLKVSYINLYCLWLLLALTILLSLILYVRKMF  571
PvPMV   LGLTFFKNRQVIFDIQKNRIGFVDANCPSHPTHTRPRTYNEYKRKDNIFLKIPFFYLYSLFVVFALSVLLSLVFYVRRLY  519
PbPMV   LGLSFFKNKQIIFDLKNNKIGFIESNCPSNPINTRPRTFNEYNIKENHLFKQSYFSLYAFSIIIALTFILYIILYIKKFI
PfPMI   LGDPFMRKYFTVFDYDNHTVGFALAKKKL---------------------------------------------------

PfPMV   YMDYFPLSDQNKSPIQEST  590
PvPMV   HMEYSPLPSEGKAPADA--  536
PbPMV   CSYYNPLHG----------
PfPMI   -------------------
```

**Supplemental Figure 3 –** Sequence alignment of the reference sequences for *P. falciparum*, *P. vivax*, and *P. berghei* PM V, as well as *P. falciparum* plasmepsin I (PfPMI). Alignments were performed using Clustal Omega. Residues shared among the PM V sequences are bolded. Structures of interest are highlighted.

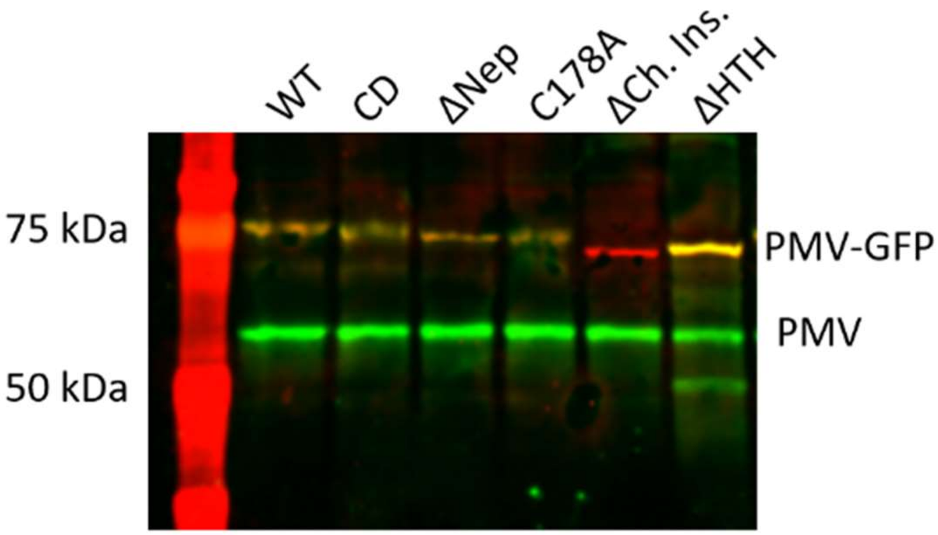


**Supplemental Figure 4 –** Western blot with anti-PMV (green) and anti-GFP (red) showing that mutants enzymes can be detected by monoclonal anti-PMV or anti-GFP, with the exception of the ΔCh. Ins. mutant, which is no longer recognized by anti-PMV.

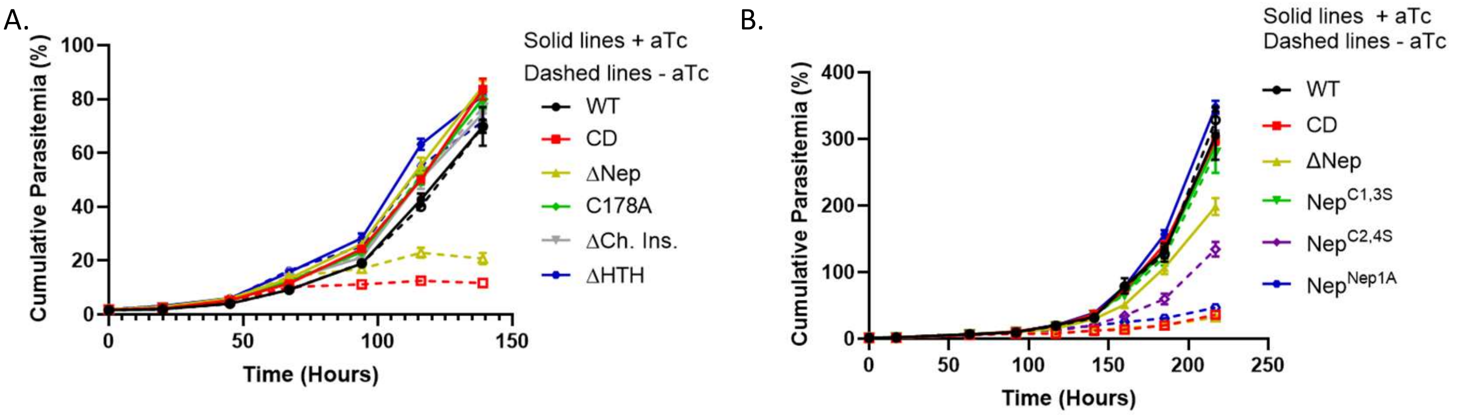


**Supplemental Figure 5 –** Full growth curves for Fig. 2D and Fig. 3D. Knockdown was performed as described above and parasite growth measured daily by flow cytometry. All parasites were subcultured 1:2 any time a well's parasitemias grew above 5%. Reads were then multiplied appropriately to calculate "cumulative parasitemias".

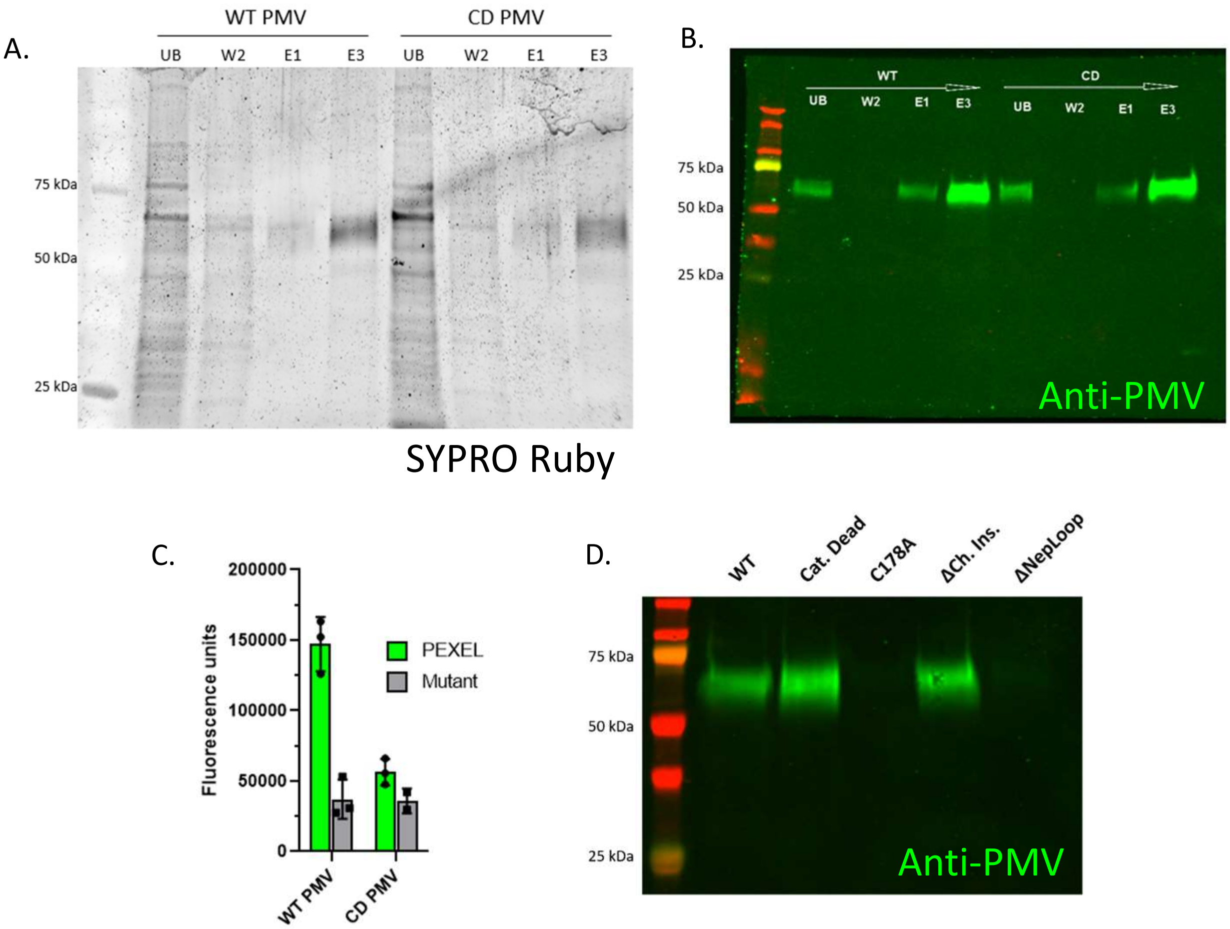


**Supplemental Figure 6 – Inconsistent expression of PM V mutants in HEK293 cells** (A) SYPRO Ruby-stained gel showing purification scheme for secreted WT and CD PM V from HEK293 conditioned media. Fractions shown are unbound (UB), a wash fraction (W2), and two elution fractions (E1 and E3). (B) Western blot of the same samples probed with anti-PM V. (C) Activity assay for WT and CD PM V showing recombinant PM V to be active against a PEXEL peptide but not the mutant peptide. Assay was performed as in Fig. 5B. (D) Western blot probed with anti-PM V.

| **Name** (use) | **Sequence** |
|---|---|
| **1** (attB into EOE) | GGTCGACTCTAGAGGATCCCCGGGTACCGAGCTCGAATTCTGGTTTGTCTGGTCAACCACCGCGGTCTCAGTGGTGTACGGTACAAACCCGAATTCTGGTTTGTCTGGTCAACCACCGCGGTCTCAGTGGTGTACGGTACAAACCCGGAATTCTAGATTTAATAAATATGTTCTTATATATAATG |
| **2** (PM V FWD) | GTGCCACCTGACGTCGAGGTGGGTGGAGGGGTTTATCCATGAGAGG |
| **3** (PM V REV) | CTGCACCTGGcctaggTGTTGATTCCTGTATGGGAG |
| **4** (PM V FWD HTH) | ACGATTTTTTctcgagATGAATAATTATTTTTTAAGGAAAGAAAATTTTTTTATATTG |
| **5** (CD) | CCATCGCAAAGAATTTCTTTAATTCTAGcTACAGGTTCATCTTCGTTAAGTTTCCCGTG |
| **6** (ΔNep) | AAACATCATCTATTTTATATGAATATCTTCAATCGTATTG |
| **7** (C178A) | GTGAATATCTTCAATCGTATgctGAAGGGTCTCAAATATATGG |
| **8** (ΔCh. Ins.) | GTTATGAACCGGATTACTTCGATATTGTGTGGCAAGCTAT |
| **9** (ΔHTH) | CCAAATAAATTATTATTTAGATATTTTATGTATACATGATATGgccgccAATTTATGTATTAAAATAGTTGATGGAGTACAATGTTGG |
| **10** ($Nep^{C1,3S}$) | CAAAAACATCATCTATTTTATATaGTAATAAATCCAATTGTCCTTATGGTTTAAAAaGTGTAGGAAATAAATGTGAATATC |
| **11** ($Nep^{C2,4S}$) | CATCTATTTTATATTGTAATAAATCCAATaGTCCTTATGGTTTAAAATGTGTAGGAAATAAAaGTGAATATCTTCAATCGTATTGTG |
| **12** ($Nep^{Nep1A}$) | GAATTATTCAAAAACATCATCTATTTTATATTGCTCAAGCCAACTCTGTCAAGCCCTTTCAAGCCCGACATGCTCTAATAATTTCTGCGAATATCTTCAATCGTATTGTGAAGGG |
| **13** (Seq 1) | GGTATTCATATGGAAAAACCATATAACTTG |
| **14** (Seq 2) | CACATATTCCAGAAAATATTTATAACC |
| **15** (Seq 3) | GTGGAAAATAAAAATGACAATGTGGGAAATAAAAATGACAATG |
| **16** (Seq 4) | CGTTAAGTTTCCCGTGTAATGGTTG |
| **17** (PMV into pHL) | GCGTAGCTGAAACCGGTgtaGAGAATAAAATCGATAATGTTGG |
| **18** (PMV into pHL) | GATGGTGGTGCTTGGTACCTGACGGGCACTTGC |
| **19** (pHL CD) | CATCCCAGCGTATATCTTTGATCTTAGcTACCGGTAGCAGCTCTTTGTCTTTTCC |
| **20** (pHL ΔNep) | CTACTCAAAAACAAGTTCTATTCTTTAtgAATACCTACAAAGTTATTGTGAAGGAAG |
| **21** (pHL C178A) | GGAAATAAATGTGAATACCTACAAAGTTATgctGAAGGAAGTCAAATATATGGTTTC |
| **22** (pHL ΔCh. Ins.) | CATCGGCGGTTATGAACCCGATTACTTcgATATTGTATGGCAAGCAATAACAAGG |
| **23** (pHL ΔHTH) | GGATATTTTGTGCATACATGACATGgccgccAACTTGTGTATAAAAATAGTTGATGG |
| **24** (pHL seq 1) | GACATAGGAAAACCATCCCAGCG |
| **25** (pHL seq 2) | GGTAGCACTTTCACGCATATACCTG |

**Supplemental Table 1 – Primers used in this study**.
